# ApoE4 accelerates the condensate to amyloid transition of tau

**DOI:** 10.1101/2025.11.22.689889

**Authors:** V Vinoth Kumar, Kanchan Garai

## Abstract

Aberrant liquid to solid transition (LST) of tau is a pathological hallmark of several tauopathies, most notably Alzheimer’s disease (AD). ApoE4 is a major genetic risk factor for AD and has been shown to exacerbate tau pathology, although the underlying mechanismremains unclear. Herewe investigate whether apoE isoforms can influence amyloid aggregation of tau occurring via the liquid–liquid phase separation (LLPS) pathway by monitoring individual tau condensates in real time using total internal reflection fluorescence microscopy (TIRFM) and confocal microscopy. We find that interaction of apoE with tau condensates is dependent on the lipidation status of apoE. Lipid-free apoE, even at 1:1000 substoichiometric concentrations, arrests growth but promotes ageing of the liquid condensates of tau. The strong effect of apoE is accompanied by its preferential accumulation inside the condensates, lowering of the surface tension and promoting amyloid nucleation. Lipidation of apoE2 and apoE3 diminished its amyloid promoting effects very strongly. However, apoE4 even in the lipidated forms accelerated the liquid to amyloid conversion of the tau condensates considerably. Furthermore, the amyloidogenic effect of apoE4 is correlated with its higher delipidation tendency compared to apoE2 and apoE3. Together, our study suggests that lipid-free apoE promotes tau aggregation via dual mechanisms by increasing condensate surface to volume ratio and by facilitating amyloid nucleation possibly at the condensate interfaces. We hypothesize that enhancing apoE4 lipidation in vivo presents a potential strategy to counteract its pathogenic effects on tau aggregation.

**Highlights:**

- All three lipid-free apoE isoforms strongly affect LLPS of tau even at a sub-stoichiometric (1:1000) ratio.
- ApoE lowers the surface tension, reduces the size and increases the number of the condensates.
- ApoE isoforms accelerate LST of the tau condensates.
- Lipidation reduces the effects of apoE, strongly in apoE2 and apoE3 but less so in apoE4.
- ApoE4 undergoes spontaneous delipidation faster than apoE3 and apoE2.

## Introduction

ApoE4 is the strongest genetic risk factor for Alzheimer’s disease^1,2^. However, the molecular mechanism by which apoE4 contributes to AD remains inconclusive. Extensive research performed over the past three decades have suggested multiple pathways for the detrimental actions of apoE4. These pathways include reduced clearance and enhanced aggregation of Aβ^3–5^, increased tau pathology^6,7^, impaired lipid metabolism^8,9^ and synaptic defects^10^, mitochondrial dysfunction^11,12^ and exacerbated neuroinflammation^13,14^. While the major volume of AD research has focused on Aβ induced pathology, mounting evidences suggest that aggregation of tau correlates more strongly with synaptic loss and cognitive decline in AD patients^15,16^.

Recent evidence from human^17,18^ and tau transgenics mice model studies^6,19^ highlights the important role of apoE4 in exacerbating tau pathology^7^. For example, in tau transgenic mice models, apoE4 has been shown to promote tau phosphorylation and aggregation, suggesting a direct role of apoE in driving tau pathology^20^. Interestingly ApoE-R136S a protective variant of apoE is shown to protect against apoE4-driven tau pathology in a mice model^21^. Moreover, human apoE and tau are observed to co-immunoprecipitate from mice brain lysate^22^, and also known to colocalize in AD brains^23,24^. However, molecular mechanism of the apoE-tau interaction is largely unexplored. Since apoE4 is the most well-established risk factor for AD^1,25–28^, understanding the mechanism of its interaction with tau can shed light on unravelling its role in exacerbating tau pathology in an isoform dependent manner.

Recent studies suggest that aggregation of tau in vivo proceeds via liquid-liquid phase separation (LLPS) ^29–33^. In this pathway soluble tau undergoes LLPS forming liquid condensates. Since tau is a highly soluble protein, condensation of tau is believed to require hyperphosphorylation^34,35^. In vitro, tau exhibits LLPS in presence of polyelectrolyte cofactors such as RNA^34,35^. Growing evidence suggests that liquid tau condensates function as hubs for microtubule nucleation playing crucial roles in stabilization of axons^36^. It is also found to regulate synaptic vesicle positioning in an activity-dependent manner^37^. However, the liquid condensates of tau are metastable. Hence, it can undergo liquid-to-solid transition (LST) via aggregation into amyloid fibrils^30,38–40^. Maturation of the condensates leads to irreversibility eventually causing impairment of the physiological functions of tau^41^.

Given the critical role of apoE4 in AD pathogenies, we examined the isoform-specific effects of apoE on tau LLPS and LST to better understand its contribution to tau pathology. Using real-time monitoring with total internal reflection fluorescence (TIRF) and confocal microscopy, we found that in the lipid-free state, all apoE isoforms strongly influence LLPS and accelerate tau amyloid aggregation. Lipidation nearly abolished the amyloid-promoting activity of apoE2 and apoE3. In contrast, apoE4 substantially retained amyloid-promoting activity even when used in the lipidated form. Further investigations revealed that the amyloidogenic effect of apoE4 stems from its tendency to undergo spontaneous delipidation. Thus, enhancing apoE4 lipidation in vivo may represent a promising therapeutic strategy to mitigate its role in tau-associated AD pathology.

## Results

### Effect of ApoE on the LLPS of tau condensates

First, we examined the effect of apoE on the LLPS of tau using all the three isoforms of apoE, *viz*, apoE2, apoE3, and apoE4 in the purified lipid-free forms. Condensates were formed by mixing 20µM P301L tau, 40 µg/ml polyU RNA in the presence of different concentrations of apoE (0 – 200 nM). We have used the P301L mutant of tau since it undergoes LLPS and subsequent ageing to amyloid fibrils in less than 24 hrs making it a tractable system^42,43^. The condensates were observed in the optical microscope by adding 5nM TMR-labeled tau in the solutions. In the lipid-free state, all the three isoforms strongly affected condensate growth. For example, the condensates are found to be smaller in size and higher in numbers in presence of apoE *(Fig 1A, B and C)*. The effect is quite dramatic even in presence of highly sub-stoichiometric concentrations (1:500 mol/mol) of apoE. For example, in presence of 200nM apoE at t=5hr, the average size reduced from 6.5µm (control) to 3.5µm and the average number of condensates per field of view (FOV ≈ 4050 µm^2^) increased from 120 (control) to 250. Expectedly, the effect is dependent on the concentrations of apoE. We also monitored the time evolution of the growth of the condensates by following both their sizes and the numbers. In absence of apoE, the number of condensates first increased with time then decreased. The decrease in number occurred due to fusion of the smaller condensates into larger condensates, consistent with the liquid nature of the condensates. However, addition of even 50 nM apoE slowed down/prevented the fusion of the tau condensates eliminating the later phase of the condensate growth *(Fig 1D).* These results indicate that lipid-free apoE strongly affects LLPS of the tau condensates. However, all the isoforms exhibited nearly identical effects (*Fig 1A-C*).

**Figure 1.**
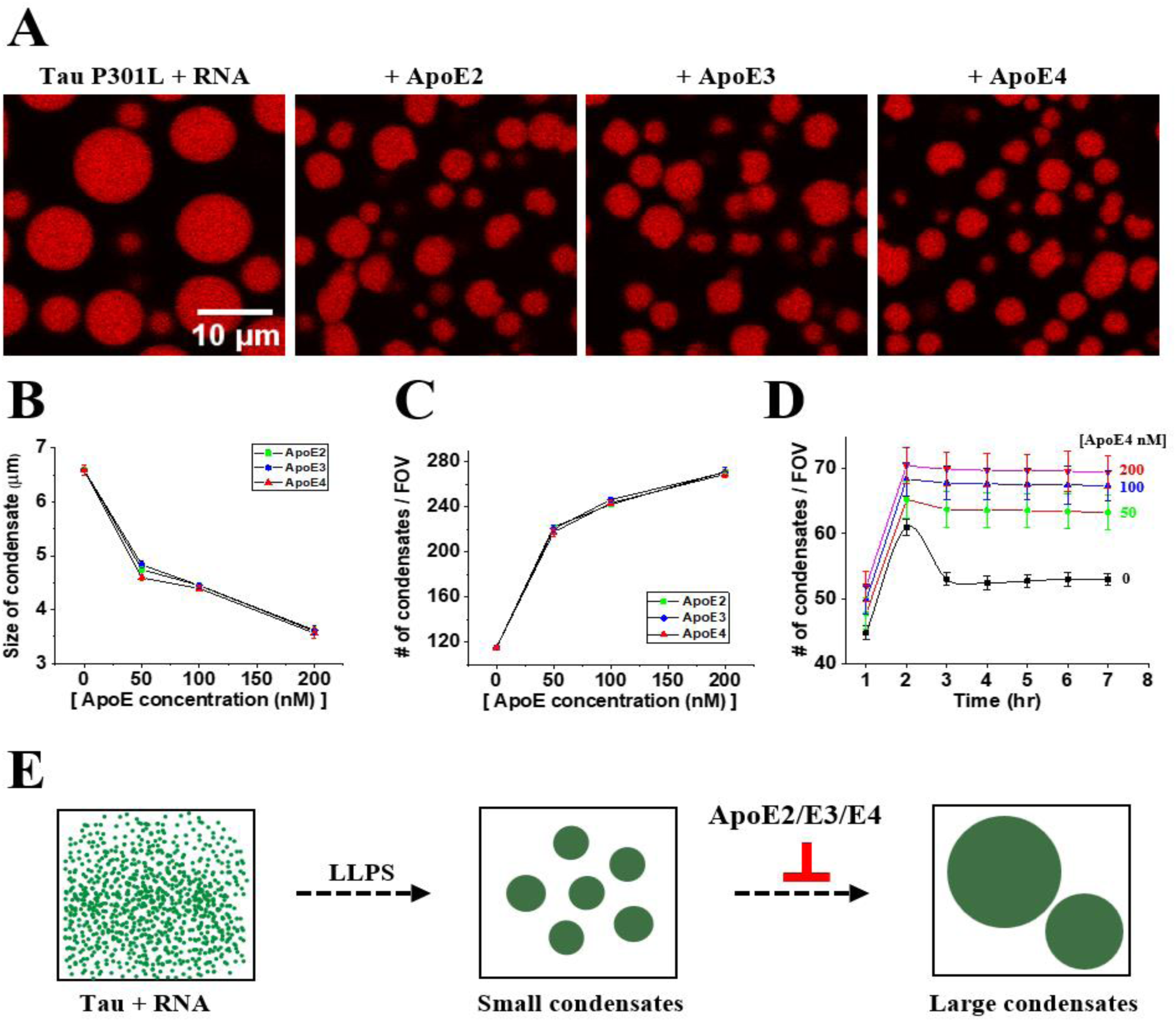
ApoE isoforms affects LLPS of tau. (A) Representative confocal microscopy images of tau condensates (20µM P301L tau, 40µg/ml polyU RNA and 5nM TMR-labeled tau) at 5hr formed in presence of no apoE or 200 nM of lipid-free apoE2, apoE3 or apoE4. (B) and (C) Size and number density distribution (Field of view, FOV = 4050µm^2^) respectively for tau condensates in absence and presence of the lipid-free apoE isoforms. (D) Time evolution of the number of the condensates per FOV (FOV = 784µm^2^). (E) A schematic summarizing the above observations that lipid-free apoE isoforms inhibits growth of the tau condensates.

### Effects of ApoE on the viscoelasticity of the tau condensates

Then we examined how apoE may alter the viscoelastic properties of the tau condensates. To examine viscosity, we performed fluorescence recovery after photobleaching (FRAP) measurements on the condensates prepared by mixing 25 nM Alexa488-labeled tau, 20 µM unlabelled tau and 40 µg/ml polyU in absence and presence of 160 nM apoE *(Fig 2A).* The FRAP kinetic data are fit using single exponential functions *(Fig 2B)* to estimate the half-times (t_1/2_), from which diffusion coefficients (*D*) were calculated using the Soumpasis equation^44^. Viscosity (η) was then estimated using the Stokes–Einstein relation, D = *k_B_T/6πηR_h_*, where *R_h_* (≈ 1.8 nm) is the hydrodynamic radius of the tau molecules *(Fig 2C)*. The viscosity of the condensates measured both in absence and presence of apoE were found to be ∼1.8 Pa·s, which is about 1800 times higher than water (*η_w_ =* 1 mPa.s). Similar value of viscosity has been reported earlier^42^. Hence, apoE does not seem to affect the viscosity of the condensates. Since apoE is present in very low substoichiometric concentrations no appreciable effect on a bulk property such as viscosity of the condensates can be expected.

**Figure 2.**
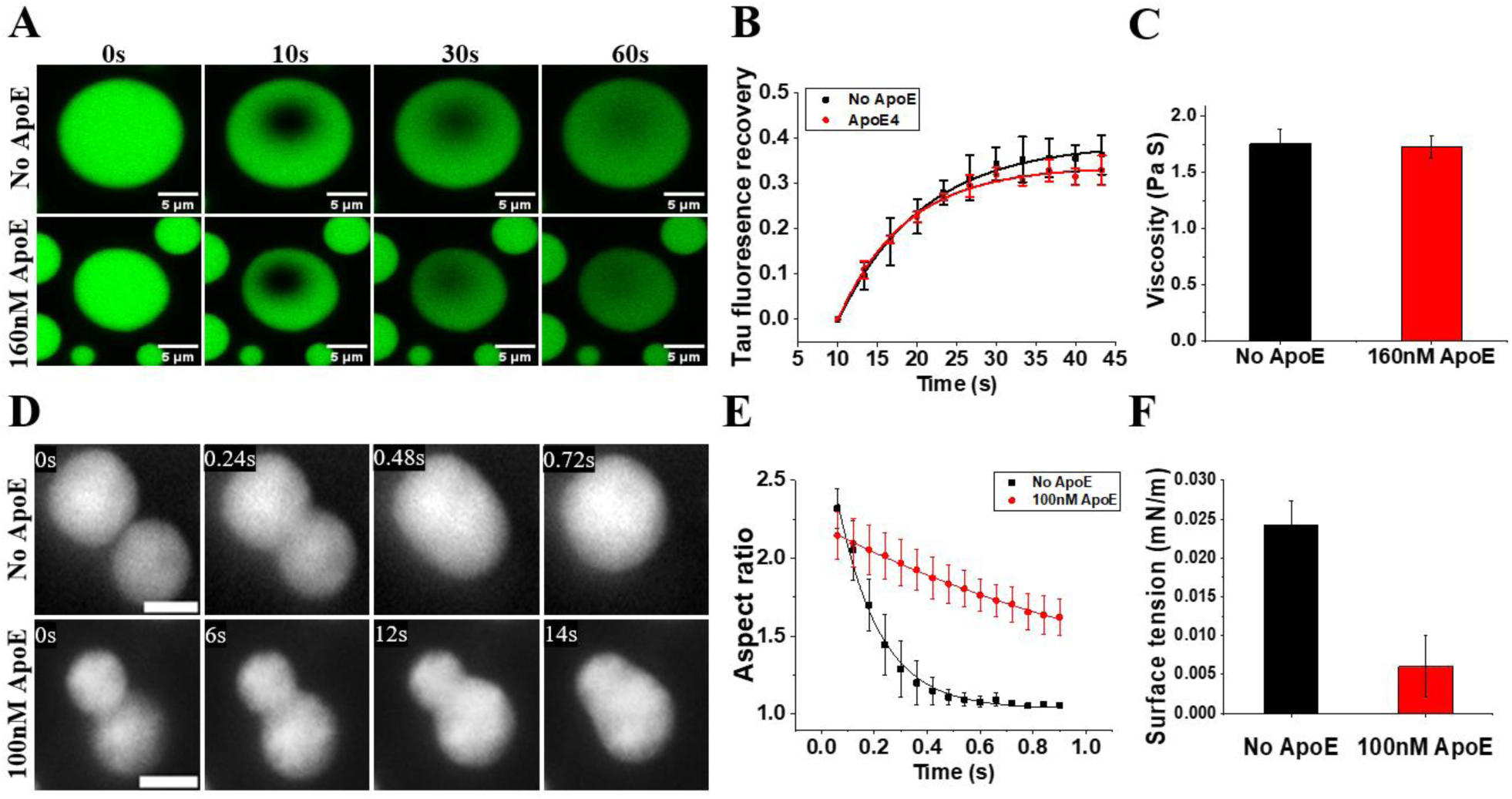
Effect of apoE on viscosity and surface tension of the tau condensates. (A) Representative confocal images showing FRAP recovery in absence and presence apoE. (B) FRAP data with fits. (C) Viscosities estimated from the FRAP kinetics. The condensates are about 1800 more viscous than water. (D) Representative images showing fu sion of the tau condensates in absence and presence of apoE. Scale 1µm (E) Fusion kinetics, monitored by the time evolution of the aspect ratio of the condensates, with fits. (G) Surface tension estimated from the fusion kinetics data. Clearly, viscosity remains unchanged, but the surface tension is reduced in presence of apoE.

Then we assessed whether apoE can alter the surface tension of the condensates. We monitored fusion dynamics of the condensates, which reflects an interplay between viscosity and surface tension. *Fig 2D* shows a few snapshots of a pair of condensates undergoing spontaneous fusion. In absence of apoE, fusion events were completed within ∼1s, with an average fusion time (τ_fusion_) ∼0.7s. Whereas in samples containing 100nM apoE, the condensates fused incompletely even after 14s, with an average τ_fusion_ *∼7.6*s. The aspect ratio relaxation kinetics were fitted to an exponential decay function to obtain the τ_fusion_ (*Fig 2E)*. The surface tension (γ) of the condensates was estimated using τ_fusion_ = (η/γ)·*l*, where η is the viscosity, γ is the surface tension and *l* is the characteristic length of the condensates. The condensates prepared in absence of apoE exhibited a γ of ∼0.025 mN/m, which is similar to that has been reported for tau biomolecular condensates^42^. However, in presence of 100 nM apoE the value of γ reduced to ∼0.005 mN/m (*Fig 2F)*. Thus, apoE reduced the surface tension of the tau condensates by about 5-fold. This is consistent with many small condensates being observed in presence of apoE. Earlier studies have shown that condensate interfaces act as sites for nucleation of amyloid aggregation^42,45,46^. Hence, apoE may enhance amyloid aggregation of tau by increasing the total interface of the condensates.

### Effect of apoE on the ageing of the tau condensates

Next, we examined whether apoE influences the ageing of tau condensates. The ageing or the liquid-to-solid transition (LST) the condensates of tau is associated with amyloid aggregation, which is a defining feature of several tauopathies ^29–32^. The ageing can be monitored by assessing the spatial inhomogeneity of the fluorescence intensity and the deformation of the shape of the condensates. At early time points, expectedly, the condensates exhibit spherical morphology and a unimodal Gaussian -like fluorescence intensity distribution. However, the intensity distribution grows more inhomogeneous, and the shape deviates to aspherical with progression of time indicating ageing (*Fig 3 A and D*). The ageing process is found to be faster in the presence of apoE. For example, the fluorescenceinhomogeneity inside thecondensates measured using coefficient of variation (defined as Standard Deviation/Mean) increased from 0.07 ± 0.02 (6hr) to 0.78 ± 0.06 in presence of 160nM apoE (25hr), while in absence of apoE the increase was minimal from 0.07 ± 0.01 to 0.08 ± 0.01 (25hr) *(Fig 3B and C)*. Similarly, the circularity index or roundness of the condensates dropped in presence of apoE (*Fig 3 E*). For example, at 25hr the circularity index dropped from 0.93 ± 0.04 (expected index =1 for a perfect circle) to 0.57 ± 0.06 in presence 160nM apoE. The aspect ratio also increased from 1.03 ± 0.04 (expected for a perfect circle) to 1.7 ± 0.23 (160nM apoE) in 25hr implying apoE mediated LST of the condensates (*Fig 3F*). Expectedly, the effect of apoE on ageing is concentration dependent. Notably, even 20 nM apoE considerably accelerated the ageing of the condensates prepared with 20 µM tau (ApoE: Tau 1:1000). Together, these data demonstrate that apoE accelerates tau ageing or LST in a concentration-dependent manner.

**Figure 3.**
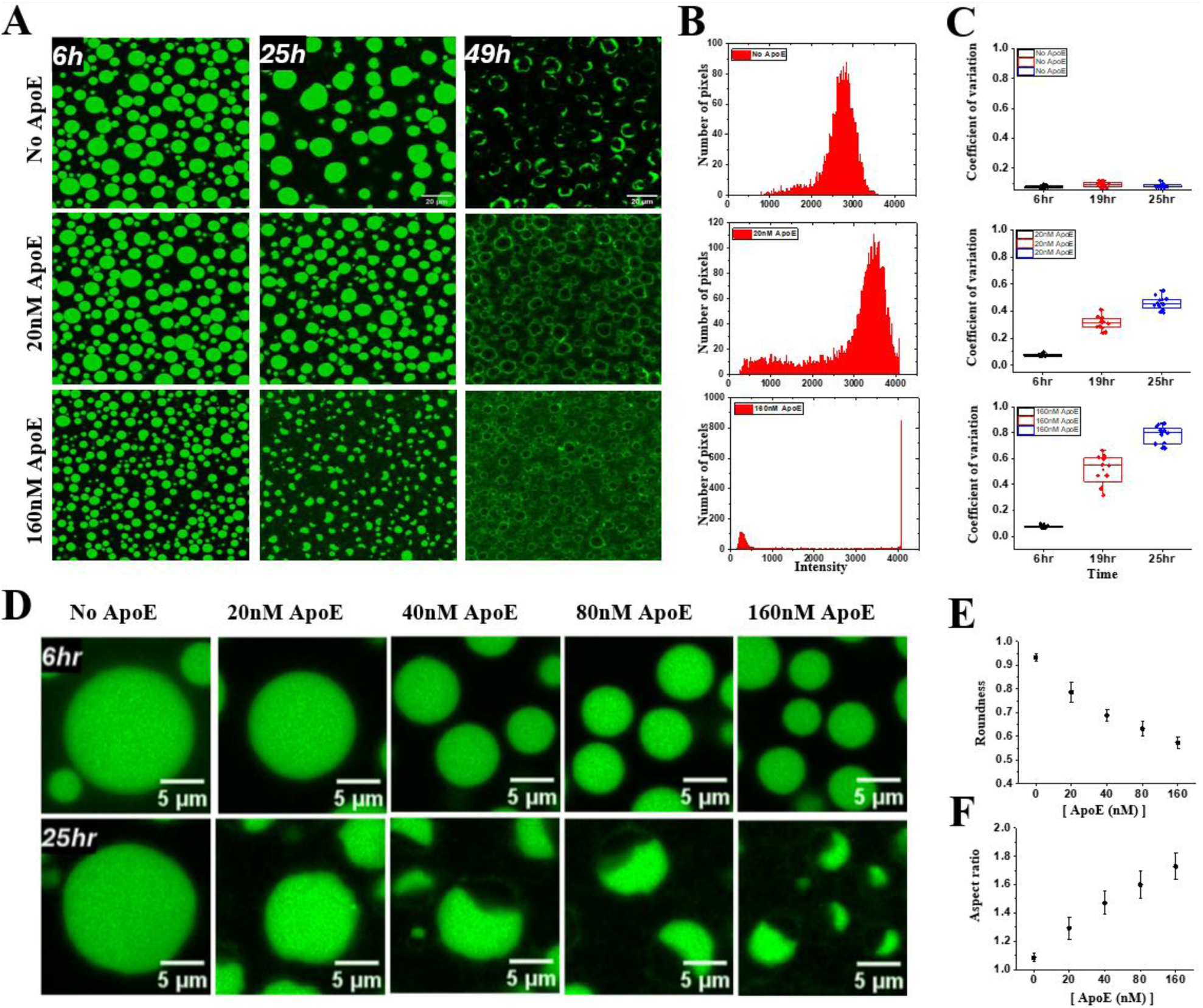
Effect of apoE on ageing of the tau condensates. (A) Representative confocal images of tau condensates prepared using 20µM P301L tau, 40µg/ml polyU RNA and 5nM alexa-488 in absence and presence of apoE4 at 6, 25 and 49hr. (B) Tau fluorescence intensity histograms of the condensates evolve from a unimodal to bimodal distribution due to liquid to solid transition at 25hr. (D) The shape of the condensates deviates from spherical to non-spherical forms upon ageing, monitored using (C) coefficient of variation of tau fluorescence intensity distribution, (E) Roundedness, and (F) Aspect ratio. Clearly, apoE accelerates ageing of the condensates.

### Effect of the ApoE on the amyloid aggregation of the tau condensates

Then we examined if apoE mediated acceleration of ageing of the condensates is correlated with the amyloid aggregation of tau. Amyloid aggregation is monitored using the fluorescence of thioflavin T and by examining the morphology in scanning electron microscopy (SEM). *Fig 4A* shows representative confocal images of the condensates formed in absence and presence of the apoE isoforms at t = 5h. The condensates, particularly the interfaces appear brighter in presence of apoE, indicating promotion of amyloid aggregation. The effect is quite dramatic in presence of 200nM apoE (apoE: tau = 1:100). For example, a 3 -fold increase of ThT fluorescence at the interface was observed in presence of 200nM apoE compared to the control. Therefore, consistent with the previous observations even a substoichiometric concentration of apoE influences amyloid aggregation of tau dramatically (*Fig 4B*). Bright fluorescenceof the condensateinterfaces is consistent with promotion of nucleation at the interface reported earlier^42,45,46^. We have also performed bulk aggregation kinetics experiments using a plate-reader to quantitate the effect of apoE on the half-time (t_1/2_) and the maximal ThT fluorescence intensity of tau aggregation (*Fig 4B).* Presence of ApoE increased the total ThT fluorescence approximately by 2-fold and the half-time (t_1/2_) of aggregation decreased from 12.5hr to 6hr (*Fig 4C* and *4D)*. Therefore, all the three isoforms of apoE promote amyloid aggregation of tau consistent with its promoting effects on the liquid to solid transition of the condensates. To investigate the morphology of the condensate and the fibrils at high resolution we performed SEM after 24hr of the sample preparation *(Fig 4E).* The SEM images show many fibrils growing inside and outside of the condensates in both cases. However, in presence of apoE the condensates exhibit a distinct morphology with the interfaces appearing as a dense and rigidified shells. Therefore, these images suggest that apoE strongly promotes maturation at the condensates interface.

**Figure 4.**
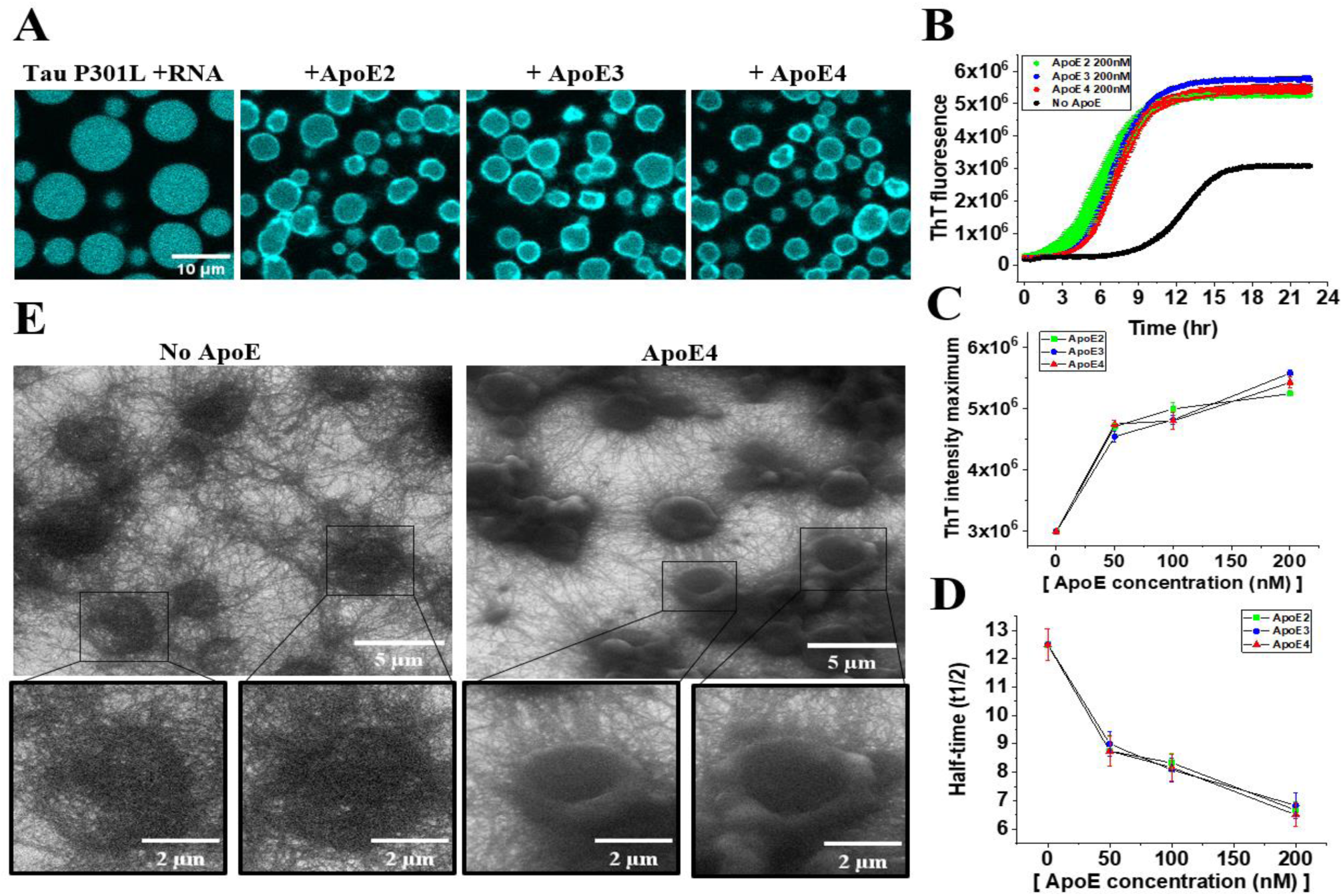
Effect of ApoE on amyloid aggregation of the tau condensates. (A) Confocal images using ThT fluorescence of tau condensates in absence and presence of 200nM apoE isoforms. Bright interfaces are observed in presence of apoE. (B) Bulk ThT fluorescence kinetics, (C) Maximum ThT intensity (at t = ∞) and (D) t _1/2_ of the aggregation kinetics in absence and presence of the apoE isoforms. (E) SEM micrograph showing the effects of 200nM apoE on the morphology of the condensates. All the figures indicate apo E accelerates amyloid aggregation of tau.

### Localization of ApoE in the condensates

Next, we examined how apoE colocalizes with tau and polyU within the condensates. In these experiments apoE is labeled with alexa-647, tau is labeled with alexa-488 and polyU is labelled with hexachlorofluorescein. First, we measured the partitioning propensities of apoE, tau and polyU inside the condensates compared to the dilute phase. All the three components were found to concentrate inside the condensates. The ratio of the concentration of apoE inside and outside of the condensates is about 300-fold for 40nM labelled lipid-free apoE. Similar fold increase of fluorescence was observed for the fluorescently labelled tau and polyU also. Thus, apoE is sequestered within the condensates with high affinity (*Fig 5A and B*). Fig 5C shows that at early time (t ∼ 2h) apoE, tau and polyU all are uniformly distributed within the milieu of the condensates, indicating fluid nature of the condensates. However, with progression of time (t ∼ 12h) distribution of apoE is shifted towards the interface (Fig 5C). Similar shift is observed in case of tau and polyU as well. Importantly, higher accumulation of the macromolecules is coincident with higher ThT fluorescence at the interface. Fold increase of the fluorescence at the interface versus the centre of the condensates are about 2.0, 3.0, 3.0 and 2.4 for apoE, tau, poly and ThT respectively. The same trend is revealed in the three-dimensional surface plot of intensity of ThT. Moreover, the surface plots highlight several bright peaks, which could be the nucleation hotspots *(Fig 5E).* ThT intensity (interface / centre) ratio clearly increases with time and concentration of apoE highlighting the apoE dependent amyloid aggregation *(Fig 5F)*. A comparative analysis of ThT and apoE fluorescence intensities indicates that localized increase in apoE concentration is closely associated with corresponding increases in ThT signal intensity. Collectively, these findings demonstrate that apoE preferentially accumulates within and along the interfaces of condensates. The strong spatial correlation between apoE and ThT distribution suggests that apoE facilitates nucleation processes and promotes amyloid conversion.

**Figure 5.**
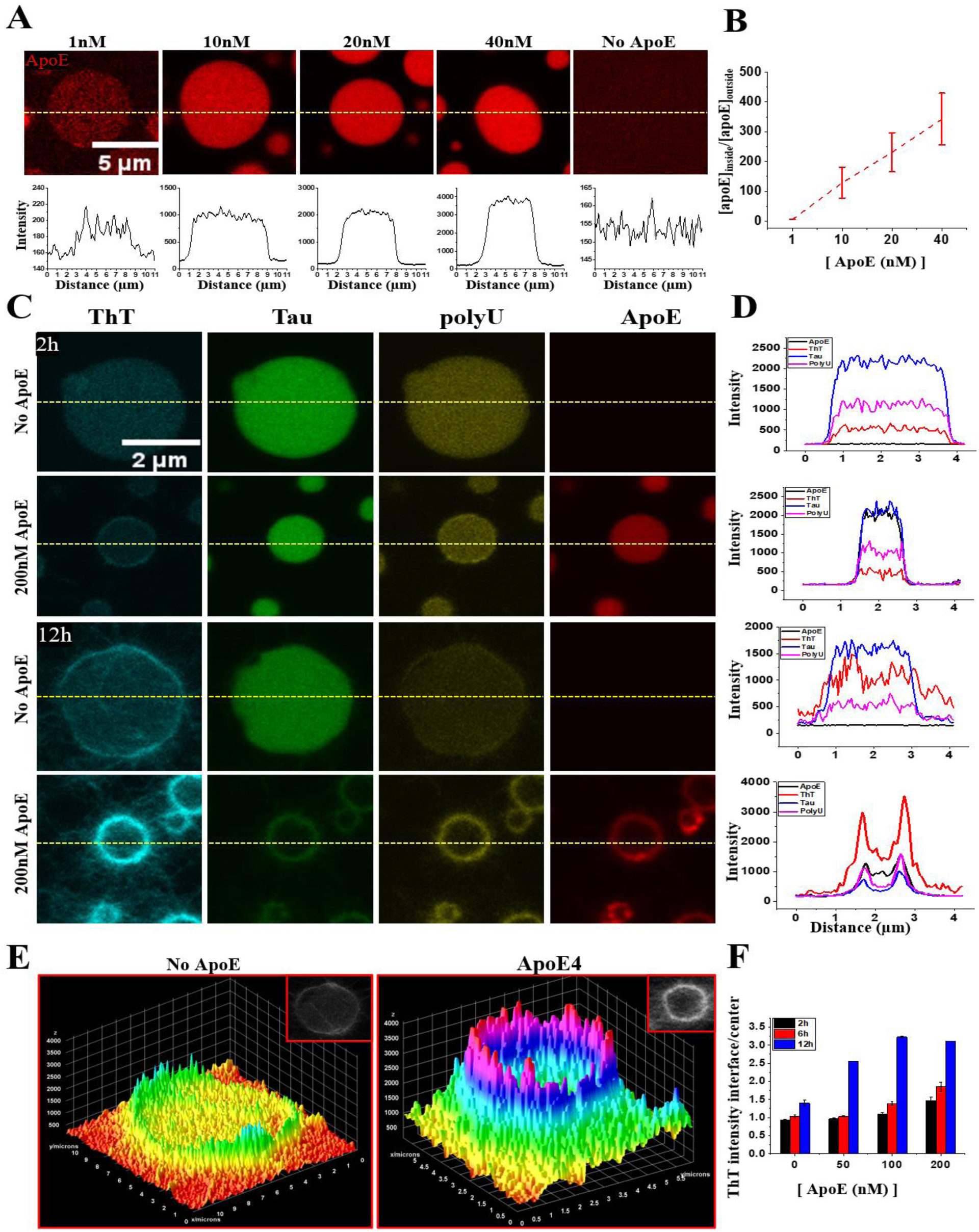
Localization of apoE in the tau condensates. (A) Confocal images of TMR-labeled apoE4 in the tau condensates at different apoE concentrations. The intensity line profiles below show strong sequestration of apoE inside the condensates. (B) Partition coefficient of apoE-TMR fluorescence (*F_inside_/F_outside_*) calculated from the confocal images. (C) Confocal images of the tau condensates imaged using ThT, alexa −488-tau, hexachlorofluorescein -polyU and Alexa-647-apoE channels at 2 and 12 hr. (D) Line profile of ThT, alexa-488-tau, hex-polyU and Alexa647-apoE across the condensate length from the confocal data. Clearly, at 12hr ThT, apoE, tau and polyU strongly colocalize at condensate interface highlighting the liquid to amyloid transition. (E) 3D surface plot of ThT intensities in absence and presence of 200n M lipid-free apoE4. (F) ThT intensity ratio (*F_interface_/F_center_*) increases with time. ApoE localizes strongly with tau.

### Effects of lipidated ApoE on the LLPS and LST of tau

Next, we examined the effect of the lipidated apoE isoforms, i.e., apoE2, apoE3, and apoE4. The lipidation of apoE was performed using DMPC^47^. Condensates were formed by mixing 20µM P301L tau, 40 µg/ml polyU RNA and 5nM TMR-labeled tau in the absence and presence of various concentrations of lipidated apoE isoforms. *Figure 6* shows that the lipidated apoE also affect the LLPS and LST of tau. However, the effects are considerably smaller than the effects observed with the lipid-free-apoE proteins. For example, the average condensate diameter decreases from 6.5 µm (control) to 5.25 µm in the presence of 200 nM lipidated apoE, (*Figs 6)* whereas a reduction of 3.5 µm was observed for 200 nM lipid-free apoE. The number of condensates observed in the presence of 200 nM lipidated apoE is 160, compared to 250 droplets formed under the same concentration of lipid-free apoE. Therefore, the effects of the lipidated apoE is considerably less compared to the lipid free apoE. However, unlike the lipid-free apoE isoforms the effect of lipidated apoE is found to be isoform dependent, with apoE4 showing the strongest effects. Imaging of the condensates using ThT fluorescence at 14hr revealed that lipidated apoE4 has stronger effects than apoE2 and apoE3 (*Fig* compared to lipidated apoE2 and apoE3. The ratio of ThT fluorescence at interface versus centre for apoE2, apoE3, apoE4 are found to be 1.4, 1.4 and 3.3 respectively (*Supplementary fig 1*)

**Figure 6.**
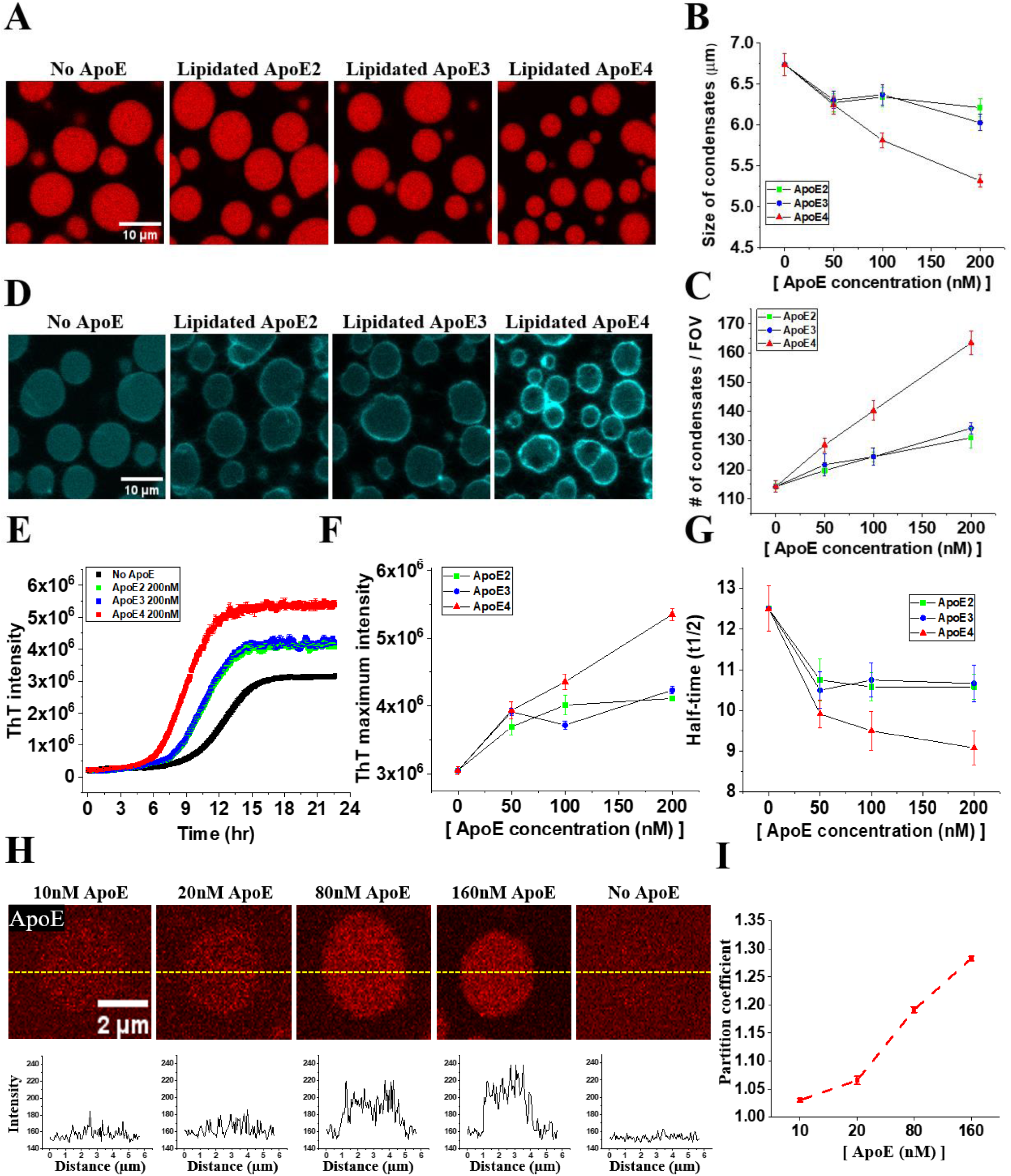
Effect of lipidated ApoE on LLPS and LST of the tau. (A) Representative confocal images of the tau condensates using fluorescence of TMR-tau(P301L) at t = 5hr in absence and presence of 200nM apoE isoforms. (B) Mean size and (C) Number of the tau condensates per FOV with apoE concentration (FOV = 4050µm^2^) (D) ThT fluorescence images of the condensates at 14hr in absence and presence of the 200nM apoE isoforms. The condensates interfaces are clearly brighter in presence of apoE4. (E) Effect of 200nM lipidated apoE isoforms on the bulk ThT fluorescence kinetics, (F) maximum ThT intensity (at t = ∞) and (G) t_1/2_ of the kinetics. (H) Uptake of lipidated TMR-apoE inside the condensates, the bottom panel shows the line profile of the intensity. (I) Ratio of TMR -apoE fluorescence inside and outside of the tau condensates. Clearly, sequestration and the effect on tau LLPS by the lipidated apoE isoforms are considerably smaller than the lipid-free apoE proteins.

Then we performed bulk aggregation kinetics measurements using ThTfluorescence in absenceand presence of the lipidated apoE isoforms *(Fig 6E).* Maximum fluorescence intensities reached ∼5 ×10^6^ units (arbitrary unit) for lipidated apoE4 versus ∼4 ×10^6^ for apoE2, apoE3 at similar concentrations and ∼3 ×10^6^ units in absence of apoE *(Fig 6H).* Therefore, lipidated apoE4 shows the highest aggregation enhancing effects. The t_1/2_ of the kinetics in presence of lipidated apoE4 (t_1/2_ ≈9–10 h) is faster, compared to that in presence of lipidated apoE2 and apoE3 (t_1/2_ ≈10.5–11 h) or in absence of any apoE (t_1/2_ ≈12 h) (*Fig 6F and G)*. Once again, we observe that lipidated apoE is less amyloid promoting than lipid-free apoE. Importantly, lipidated apoE4 exhibits the strongest effects compared to lipidated apoE2 and apoE3. Then we examined the propensity of partitioning of lipidated apoE within the condensates. We used fluorescently labelled lipidated apoE to compareits concentrationsinsidewith that of the outside the condensates *(Fig 6H).*The fluorescence of apoE inside the condensates is found to be slightly higher (about 2.0 -times for 160 nM DMPC-TMR-apoE4) than that of the outside. It may be recalled that in case of lipid-free apoE the partition coefficient was more than 300 for 40nM TMR-apoE (see Fig 5A).

Therefore, unlike lipid-free apoE, lipidated apoE partitions poorly within the condensates. This is consistent with the considerably poor effects of lipidated apoE isoforms on the LLPS and LST of tau.

### Comparing delipidation kinetics of the apoE isoforms

Strong affinity of apoE for the tau condensates in the lipid-free form but poor affinity in the lipidated form led us to hypothesize that lipidated apoE undergoes slow spontaneous delipidation (*Fig 7A*), and only the lipid-free form of apoE interacts with the condensates. We note here that the ApoE proteins are known as exchangeable lipoproteins due to their functions as lipid transporters^48–50^. Hence, *in-vivo* functions of apoE requires it to transition between lipid-poor and lipid rich forms^51–53^. Therefore, we set out to compare the apoE isoforms in terms of its delipidation propensities starting from the lipidated state. The difficulty in quantification of the delipidation of apoE is that the population of lipid-free apoE is extremely small compared to the lipidated apoE since in presence of lipids the equilibrium is strongly shifted towards the lipidated state. Therefore, we employed a novel strategy to compare the delipidation rates of apoE3 and apoE4. First, we prepared lipidated apoE3 and apoE4 using DMPC. These samples were then subjected to KBr density-gradient ultracentrifugation for 18hr and 40 hr. The lipidated apoE is expected to float up and delipidated apoE to move towards the bottom of the centrifuge tube ^54^. Then we fractionated the samples based on its height in the tube and measured the concentration of apoE in these fractions *(Fig. 7E).* The results show that in the top fractions concentrations of apoE3 is higher indicating more lipidated apoE3 than apoE4. However, in the bottom fractions apoE4 is more concentrated than apoE3 indicating more lipid-free apoE4 (*Fig 7C and D*). Moreover, with progression of time (18 hr versus 40 hr) the concentration of lipid-free apoE4 increased faster than that of apoE3 (*Fig 7D*). Therefore, apoE4 undergoes faster delipidation than apoE3. This is consistent with reports of poor lipidation status of apoE4 than apoE3 *in-vivo*^51,55^.

**Figure 7.**
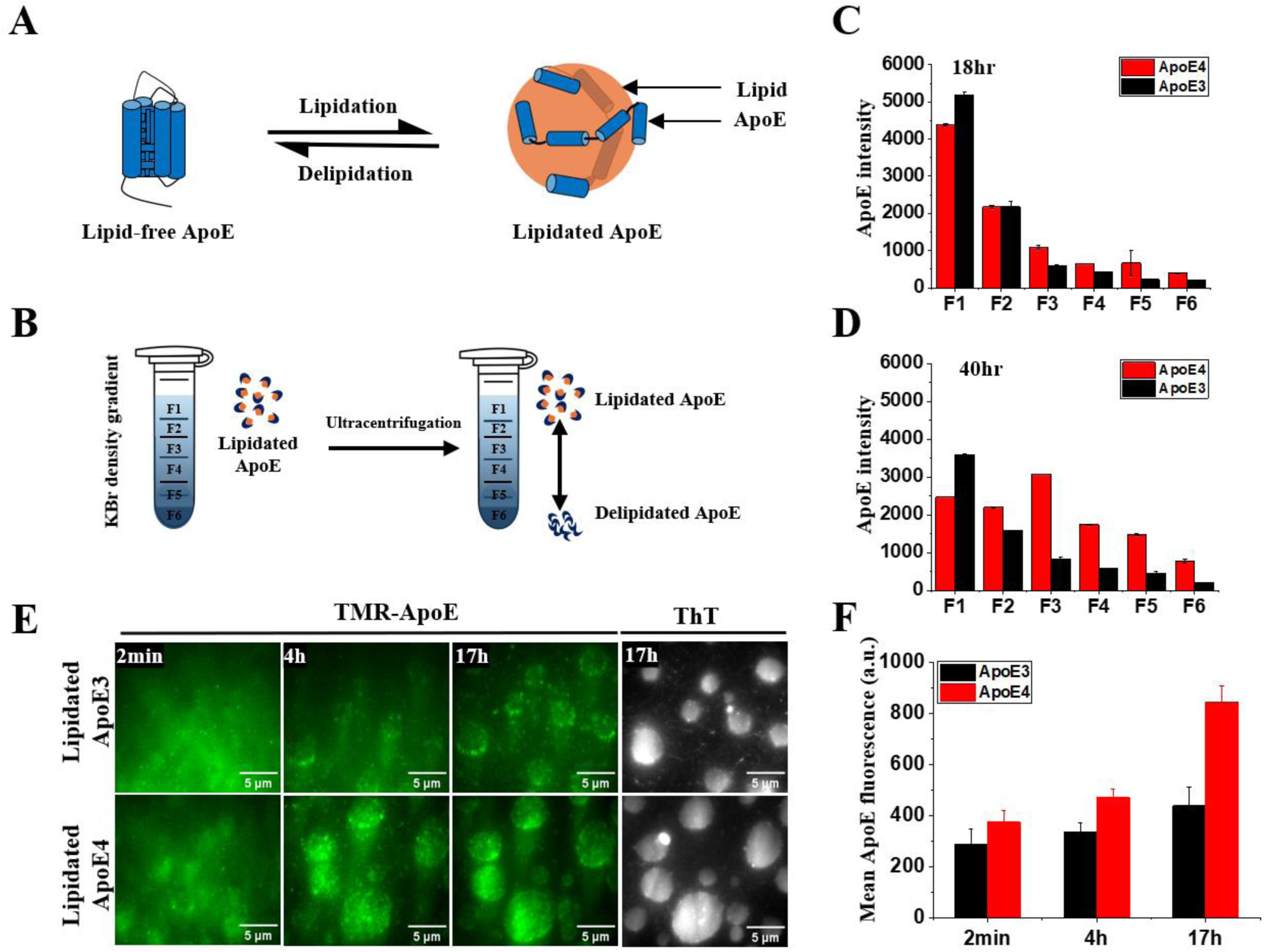
Lipidation-delipidation dynamics drives the isoform-specific effects of apoE. (A) Schematic showing apoE lipidation and delipidation dynamics. (B) Ultracentrifugation in KBr density gradient is used to monitor delipidation of apoE as a function of time. KBr densities are 1.3 and 1.1g/ml at the bottom and top of the tube respectively. (C) and (D) ApoE concentration from top to bottom monitored by intrinsic tryptophan fluorescence. (E) TIRF images showing entry of TMR-apoE3 or apoE4 inside the tau condensates 2min, 4hr and 17hr after addition in the lipidated forms. (F) Mean fluorescence intensity of TMR-apoE3 and apoE4 inside the condensate. Clearly, apoE4 exhibits higher delipidation tendency and faster sequestration within the condensates.

Next, we argued that if the sequestration of apoE inside the tau condensates is dependent on its delipidation, the concentration of apoE inside the condensates should slowly increase with time, if we start with the lipidated form of apoE. Furthermore, we expect th is kinetics to be isoform dependent (apoE4>apoE3). Hence, we prepared TMR-labeled DMPC-apoE3 and apoE4 and monitored its time dependent entry inside the condensates using fluorescence microscopy. Importantly, in this experiment we used the condensates of WT tau, which is stable in the liquid state for several days without undergoing LST (*supplementary Fig 2*). *Figure 7E* shows that fluorescence of both apoE4 and apoE3 inside the condensate increases with time. Notably, the fluorescence of TMR-apoE4 is considerably higher than TMR-apoE3 at both t = 4 and 17 hr *(Fig 7F).* Therefore, we hypothesize that delipidation of apoE is required for sequestration into the condensates, and this process is faster in case of apoE4.

## Discussion

Despite high solubility of tau in native solution conditions it undergoes irreversible amyloid aggregation, *in vivo*, which is a hallmark of the tauopathies. What triggers the aggregation of tau is still unknown. An emerging hypothesis is that biomolecular condensates of tau undergo LST leading to formation of fibrillar aggregates under pathological conditions^29,40^. However, physiological factors that promote or inhibit the transition of the soluble tau into insoluble aggregates is still unclear. Previous genome wide association studies showing association of apoE with CSF tau and p-tau independent of Aβ ^56,57^ and the linkage of apoE with the primary tauopathies^25–27^ suggest that apoE may have a role in regulating tau pathogenesis. Early studies have shown that apoE isolated from human plasma binds to recombinant tau in an isoform-specific fashion in vitro^58,59^. In addition, apoE immunoreactivity is found to colocalize with neurofibrillary tangles^23^. Overexpression of human apoE4 in neurons, but not astrocytes, has been reported to increasetau phosphorylation in mousebrains compared to apoE3^60,61^. ApoE4 is shown to significantly increase tau hyperphosphorylation, neuroinflammation and brain atrophy compared to other apoE isoforms in tauopathy mice models^6^.Importantly, selective removal of astrocytic apoE4^62^ or lowering of apoE4 level with antisense oligonucleotides^63^ has been shown to reduce tauopathy and neurodegeneration in mouse models. Tau, primarilyintracellular, is also secreted into the brain ISF^64^, where it is believed to transmit tau pathology in a prion-like manner^65^.Abundance of apoE in CSF and ISF could possibility regulate spreading of tau aggregates by direct interaction. Recent high-resolution transcriptomic data on tau accumulation and spreading in AD brains suggested apoE as a central player in the spatial spreading of tau^66^. Importantly, apoE3-R136C, a protective apoE variant, was shown to strongly bind to tau, inhibit its uptake by neurons^67^ and also previously reported to provide protections from tau mediated neurodegeneration in mice^21^. Alternatively, overexpression of low-density lipoprotein receptor (LDLR) is also shown to reduce tau-associated neurodegeneration in a mice model by reducing apoE levels^68^. Together, these studieshighlight the pathogenic rolesof apoE4 in modulatingtau phosphorylation, inducing microglial responses and neuroinflammation. However, mechanisms by which apoE4 promote tau pathologies are not clear. Since, apoE colocalizes with tau in vivo^23^ and found to bind to tau in vitro^58,67^, we asked whether apoE can directly influence the amyloid aggregation of tau.

In our study we find that all three apoE isoforms in their lipid-free statesdramatically accelerate amyloid aggregation mediated by LLPS of tau even at low substoichiometric concentrations (*ApoE: Tau = 1:1000*). Effectiveness of apoE in promoting amyloid aggregation of tau at very low substoichiometric concentrations suggests that apoE acts a pathological chaperone for tau aggregation. We think that apoE promotes the nucleation of tau aggregation, which is evident from the rapid increase of ThT fluorescence at the interface of the tau condensates. Moreover, we find that apoE lowers the surface tension of the condensates leading to stabilization of numerous smaller condensates. This leads to increase of the total surface area of the condensates. Earlier reports have suggested that the condensate interfaces act as sites for nucleation of amyloid aggregation of several proteins involved in neurodegeneration^42,45,46^. Therefore, stabilization of the smaller condensates possibly enhances the amyloid promoting role of apoE even further.

A striking observation in this paper is that lipidation of apoE reduces this interaction strongly. This is especially high for apoE2 and apoE3. Therefore, in lipidated forms of apoE2 and apoE3 are almost benign. This is consistent with the non-pathogenic role of these two isoforms of apoE. However, lipidated apoE4 exhibited considerably accelerated the LST of the tau condensates. Therefore, lipidation was not sufficient to abolish the amyloid promoting effect of apoE4. Investigation into the mechanism of this action using a novel density gradient ultracentrifugation method revealed that delipidation kinetics of apoE4 is much faster than the other isoforms. Therefore, our studies suggest that lipidated apoE4 by itself is non-pathogenic. However, faster delipidation of apoE4 leads to enhancement of its pathogenic effects. Interestingly, tau transgenic mouse models showed higher tau pathology and neurodegeneration in presence of ApoE4 compared to ApoE3 or ApoE2. Importantly, such pathological effects were rescued by inducing lipidation of ApoE using LXR agonists^69^. Thus, we propose that promoting lipidation of apoE4 in vivo can ameliorate its pathogenic effects. *Figure 8* shows a schematic model on the effect of apoE on LLPS and LST of tau condensates

**Figure 8.**
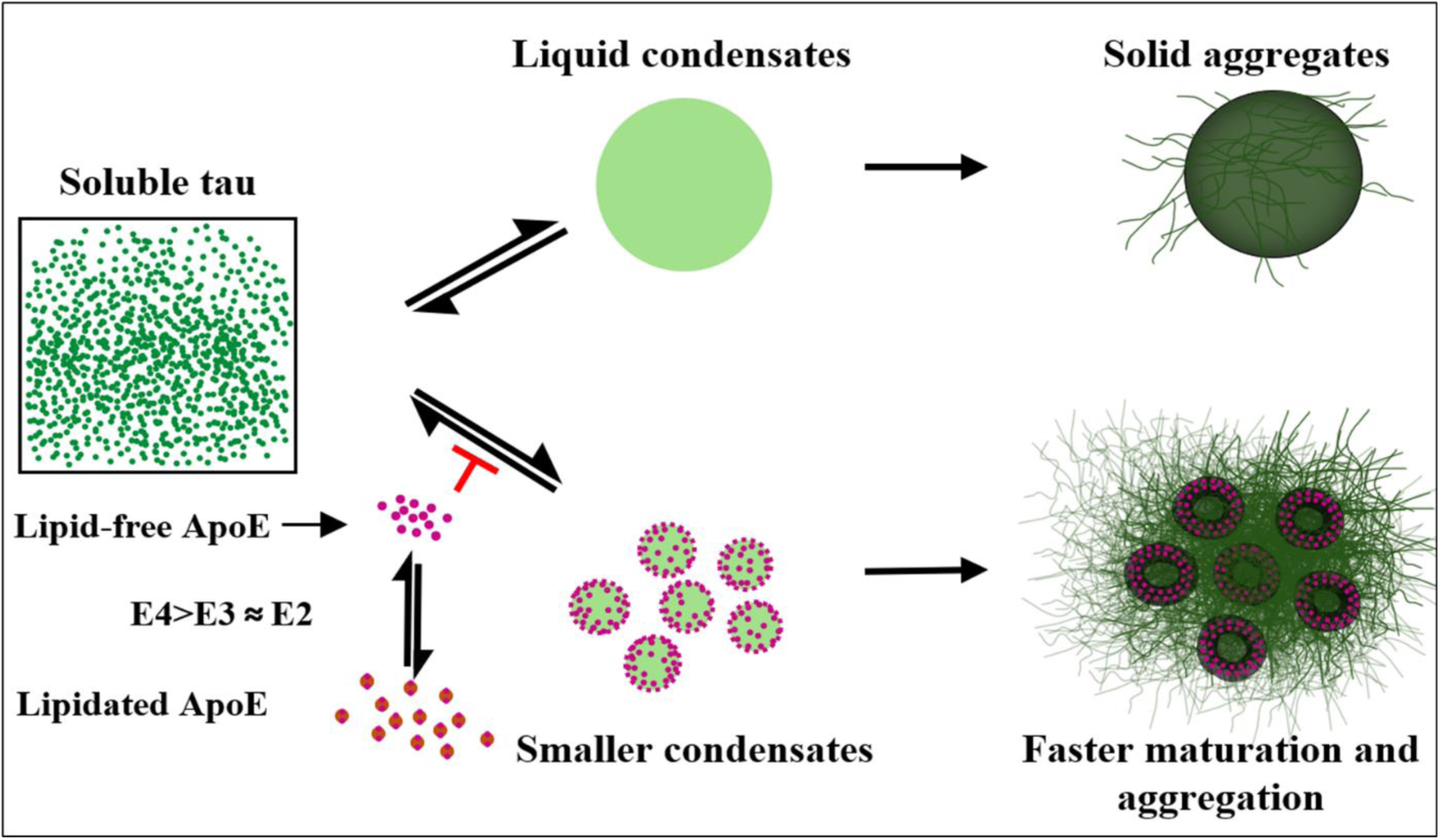
Schematic representation showing the effect of apoE on LLPS and LST of tau condensates. Soluble tau in presence of RNA undergoes reversible phase separation to form liquid condensates, which matures into solid aggregates by ageing. Increased delipidation of apoE4 results in lipid-free apoE4, which actively partitions inside the tau condensates and impairs condensate fusion by lowering its surface tension. The impaired fusion process results in smaller and more numerous condensates, thus affecting liquid–liquid phase separation (LLPS). Lipid-free apoE preferentially localizes at the condensate interface, where it promotes amyloid nucleation and facilitates the liquid-to-solid transition (LST) of tau condensates through interface-driven nucleation.

In general, condensates facilitate transient interactions to facilitate processes like transcription, RNA metabolism, and signal transduction^70^. By concentrating monomers, the condensates can accelerate reactions, at the same time the liquid nature of the condensates allows maintaining of protein homeostasis in vivo. This is crucial for the biological functions of the biomolecular condensates^37^. Since, LLPS of tau is shown to be important for its microtubule stabilizing function, irreversible conversion of the liquid condensates to solid forms by apoE4 can affect the physiological functions of tau such as nucleation and organization of the microtubule^50^ and activity-dependent synaptic vesicle recycling^47^. Thus, apoE4 may cause loss of function of the intracellular tau by promoting ageing of the tau condensates.

## Conclusion

In summary, all the isoforms of apoE affects both LLPS and accelerate LST of the tau condensates in the lipid free state. Presence of apoE, leads to increase in the total surface area of tau condensates generating more interfacial regions that possibly function as hotspots for tau nucleation and thereby accelerate the liquid-solid transition (LST) to amyloid fibrils. However, in the lipidated forms only apoE4 affected LLPS and accelerated LST of tau appreciably. ApoE4 specific effects on the tau condensates correlates with its increased propensity for spontaneous delipidation. Therefore, our study suggests enhancing lipidation of apoE4 should be pursued as strategy to mitigate its pathogenic role in AD.

## Materials and methods

All the chemicals, unless otherwise specified, were sourced from Sigma Aldrich (MO, USA). The WT-apoE4 plasmid was kindly provided by Dr. Carl Frieden from Washington University, School of Medicine, St. Louis, USA. 2N4R Tau (residues 255–441) plasmid was a kind gift from Songi Han lab (University of California, Santa Barbara). All other plasmids are derived from WT-apoE4 plasmid or 2N4R Tau plasmid by site-directed mutagenesis which was carried out by Genescript (USA).

### Protein expression and purification

#### Tau

In all experiments, the truncated variant of 2N4R tau (residues 255–441) was used. This variant includes the disease-relevant P301L mutation along with two additional mutations (C291S and C322S) to eliminate native cysteine residues. Additionally, a 6×His tag was added at the N-terminal to facilitate protein purification. The detailed protocol for protein expression and purification is described elsewhere^72^. Briefly, the protein was expressed in E. coli BL21(DE3) cells and purified using Ni-NTA chromatography following standard protocols. The purified protein was subjected to buffer exchange through dialysis in 20 mM HEPES buffer (pH 7.4) utilizing a 10 kDa dialysis bag. Purity was confirmed through SDS-PAGE and mass spectrometry analysis, and protein concentration was measured via UV-Vis absorbance at 277 nm using a molar extinction coefficient of 2980 cm⁻¹M⁻¹.

#### Tau-labelling

For fluorophore labelling, P301Ltau containing a single cysteine residue (C291S/P301L/C322) was used. Labelling was performed with Alexa488 maleimide, and the labelled protein was purified by size exclusion chromatography using a Superdex 200 column (GE Healthcare)^42^.

#### ApoE

Full-length apoE are expressed as fusion proteins with a 6×-His tag, the fusion partner thioredoxin followed by a PreScission protease cleavage site attached at the N-terminal of the apoE. ApoE proteins are expressed and purified using protocols described elsewhere^73^. The final step of the purification of the proteins is size exclusion chromatography (SEC) using a Superdex 200 column (GE Healthcare, USA) in 20mM HEPES buffer 50mM Nacl at pH 7.4 Concentration of the proteins are determined using absorbance at 280 nm. The extinction coefficients of the full-length protein ApoE is 44460 cm⁻¹M⁻¹ used for concentration measurements. The purified stock solution was used within 3 days and regularly purified before each experiment.

#### ApoE-labelling

Single cysteine mutants of apoE (e.g., apoE(W210C) are labeled with alexa647-maleimide (Anaspec, USA) or TMR maleimide using standard protocol of maleimide labeling^74^. The fluorescently labeled protein is purified by SEC using a Superdex 200 column in 20mM HEPES buffer 50mM Nacl at pH 7.4 containing 1 mM EDTA and 5 mM βMe. Purified stock solution is then aliquoted, flash frozen and stored at −80 °C.

#### ApoE-lipidation

Lipidation of apoE was performed by incubating the protein with 1,2-dimyristoyl-sn-glycero-3-phosphocholine (DMPC) liposomes in phosphate-buffered saline (PBS) at 25°C overnight, maintaining a protein-to-liposome ratio of 1:60. Following incubation, the mixture subjected to KBr density gradient ultracentrifugation at 80,000×g and 15°C overnight to separate lipidated ApoE from its lipid-free counterpart. The lipidated form was recovered from the upper, less dense layer (density = 1.1 g/mL). Finally, unbound lipids and lipidated apoE were separated by size exclusion chromatography using a Superdex 200 column (GE Healthcare, USA) in 20mM HEPES buffer 50mM Nacl at pH 7.4. Lipidation was confirmed by a characteristic blue shift in tryptophan fluorescence upon lipid interaction^54,75^. For labelled lipidated apoE, we first labelled apoE with TMR then performed the lipidation of the labelled apoE.

#### Sample preparation for the microscopy experiments

A coverslip bottom multi-well chamber (4-well or 96-well, Celvis) was treated with 10 M NaOH and rinsed with MilliQ water several times to remove contaminants^76^. P301L tau stock solution was diluted to 10 to 20uM concentration in 20 mM Hepes buffer (pH 7.4) containing 10 µM ThT, 10 mM βMe, and 40 µg/ml PolyU RNA to induce LLPS. Imaging was performed using a home-built TIRF microscope^76^, ThT fluorescence was used to identify the liquid condensates and the amyloid fibrils. The 3D Z-stack imaging and FRAP measurements were performed using a confocal microscope (Olympus FV3000) with 25nM Alexa-488-labeled P301L tau to assess the viscosity of the condensates.

#### Sample preparation for SEM morphological analysis

A small (5mmx 5mm) silicon wafer was placed in a TIRF sample chamber, containing tau and polyU. The assembly was incubated at 25°C for 24 hours to allow formation of condensates and subsequently the fibrils. After incubation, the silicon wafer was rinsed gently in Milli-Q water bath several times to remove excess buffer and any unbound materials. Then it was dried in air for overnight. The dried wafer was imaged using scanning electron microscopy (SEM) to examine the morphology of the condensates and the fibrils.

## Acknowledgement

The authors thank Dr. Yann Fichou for sharing the tau plasmid with us. We also thank Mahaprasad S.R. Sahu and Deepa.S for their help with tau and apoE expression and purification. Sincere thanks to Shravani, Akhil for their help in scanning electron microscopy and confocal imaging. This work received generous support from the Department of Atomic Energy, Government of India (project ID no. RTI 4007 to V.K.V. and K.G.), the Science and Engineering Research Board, Government of India (grant no. CRG/2020/005527 to K.G.). We thank Prof. Tuomas P. J. Knowles for valuable discussions and suggestions.

## Reference

(1) Corder, E. H.; Saunders, A. M.; Strittmatter, W. J.; Schmechel, D. E.; Gaskell, P. C.; Small, G. W.; Roses, A. D.; Haines, J. L.; Pericak-Vance, M. A. Gene Dose of Apolipoprotein E Type 4 Allele and the Risk of Alzheimer’s Disease in Late Onset Families. Science 1993, 261 (5123), 921–923. 10.1126/science.8346443.

(2) Liu, C.-C.; Kanekiyo, T.; Xu, H.; Bu, G. Apolipoprotein E and Alzheimer Disease: Risk, Mechanisms and Therapy. Nat Rev Neurol 2013, 9 (2), 106–118. 10.1038/nrneurol.2012.263.

(3) Liu, C.-C.; Zhao, N.; Fu, Y.; Wang, N.; Linares, C.; Tsai, C.-W.; Bu, G. ApoE4 Accelerates Early Seeding of Amyloid Pathology. Neuron 2017, 96 (5), 1024–1032.e3. 10.1016/j.neuron.2017.11.013.

(4) Lin, Y.-T.; Seo, J.; Gao, F.; Feldman, H. M.; Wen, H.-L.; Penney, J.; Cam, H. P.; Gjoneska, E.; Raja, W. K.; Cheng, J.; Rueda, R.; Kritskiy, O.; Abdurrob, F.; Peng, Z.; Milo, B.; Yu, C. J.; Elmsaouri, S.; Dey, D.; Ko, T.; Yankner, B. A.; Tsai, L.-H. APOE4 Causes Widespread Molecular and Cellular Alterations Associated with Alzheimer’s Disease Phenotypes in Human iPSC-Derived Brain Cell Types. Neuron 2018, 98 (6), 1141–1154.e7. 10.1016/j.neuron.2018.05.008.

(5) Kim, J.; Basak, J. M.; Holtzman, D. M. The Role of Apolipoprotein E in Alzheimer’s Disease. Neuron 2009, 63 (3), 287–303. 10.1016/j.neuron.2009.06.026.

(6) Alzheimer’s Disease Neuroimaging Initiative; Shi, Y.; Yamada, K.; Liddelow, S. A.; Smith, S. T.; Zhao, L.; Luo, W.; Tsai, R. M.; Spina, S.; Grinberg, L. T.; Rojas, J. C.; Gallardo, G.; Wang, K.; Roh, J.; Robinson, G.; Finn, M. B.; Jiang, H.; Sullivan, P. M.; Baufeld, C.; Wood, M. W.; Sutphen, C.; McCue, L.; Xiong, C.; Del-Aguila, J. L.; Morris, J. C.; Cruchaga, C.; Fagan, A. M.; Miller, B. L.; Boxer, A. L.; Seeley, W. W.; Butovsky, O.; Barres, B. A.; Paul, S. M.; Holtzman, D. M. ApoE4 Markedly Exacerbat es Tau-Mediated Neurodegeneration in a Mouse Model of Tauopathy. Nature 2017, 549 (7673), 523–527. 10.1038/nature24016.

(7) Lennol, M. P.; Bordier, C.; Kamelher, L.; Ulrich, J. D.; Holtzman, D. M.; Gratuze, M. ApoE-Calypse Tau: ApoE–Tau Synergy in Alzheimer’s Disease. Journal of Experimental Medicine 2025, 222 (10), e20250965. 10.1084/jem.20250965.

(8) Haney, M. S.; Pálovics, R.; Munson, C. N.; Long, C.; Johansson, P. K.; Yip, O.; Dong, W.; Rawat, E.; West, E.; Schlachetzki, J. C. M.; Tsai, A.; Guldner, I. H.; Lamichhane, B. S.; Smith, A.; Schaum, N.; Calcuttawala, K.; Shin, A.; Wang, Y.-H.; Wang, C.; Koutsodendris, N.; Serrano, G. E.; Beach, T. G.; Reiman, E. M.; Glass, C. K.; Abu-Remaileh, M.; Enejder, A.; Huang, Y.; Wyss-Coray, T. APOE4/4 Is Linked to Damaging Lipid Droplets in Alzheimer’s Disease Microglia. Nature 2024, 628 (8006), 154–161. 10.1038/s41586-024-07185-7.

(9) Holtzman, D. M.; Herz, J.; Bu, G. Apolipoprotein E and Apolipoprotein E Receptors: Normal Biology and Roles in Alzheimer Disease. Cold Spring Harbor Perspectives in Medicine 2012, 2 (3), a006312–a006312. 10.1101/cshperspect.a006312.

(10) Chen, Y.; Durakoglugil, M. S.; Xian, X.; Herz, J. ApoE4 Reduces Glutamate Receptor Function and Synaptic Plasticity by Selectively Impairing ApoE Receptor Recycling. Proc. Natl. Acad. Sci. U.S.A. 2010, 107 (26), 12011–12016. 10.1073/pnas.0914984107.

(11) Schmukler, E.; Solomon, S.; Simonovitch, S.; Goldshmit, Y.; Wolfson, E.; Michaelson, D. M.; Pinkas-Kramarski, R. Altered Mitochondrial Dynamics and Function in APOE4 -Expressing Astrocytes. Cell Death Dis 2020, 11 (7), 578. 10.1038/s41419-020-02776-4.

(12) Yin, J.; Reiman, E. M.; Beach, T. G.; Serrano, G. E.; Sabbagh, M. N.; Nielsen, M.; Caselli, R. J.; Shi, J. Effect of ApoE Isoforms on Mitochondria in Alzheimer Disease. Neurology 2020, 94 (23). 10.1212/WNL.0000000000009582.

(13) Chen, Y.; Jin, H.; Chen, J.; Li, J.; Găman, M.-A.; Zou, Z. The Multifaceted Roles of Apolipoprotein E4 in Alzheimer’s Disease Pathology and Potential Therapeutic Strategies. Cell Death Discov. 2025, 11 (1), 312. 10.1038/s41420-025-02600-y.

(14) Parhizkar, S.; Holtzman, D. M. APOE Mediated Neuroinflammation and Neurodegeneration in Alzheimer’s Disease. Seminars in Immunology 2022, 59, 101594. 10.1016/j.smim.2022.101594.

(15) Josephs, K. A.; Whitwell, J. L.; Ahmed, Z.; Shiung, M. M.; Weigand, S. D.; Knopman, D. S.; Boeve, B. F.; Parisi, J. E.; Petersen, R. C.; Dickson, D. W.; Jack, C. R. Β-amyloid Burden Is Not Associated with Rates of Brain Atrophy. Annals of Neurology 2008, 63 (2), 204–212. 10.1002/ana.21223.

(16) La Joie, R.; Visani, A. V.; Baker, S. L.; Brown, J. A.; Bourakova, V.; Cha, J.; Chaudhary, K.; Edwards, L.; Iaccarino, L.; Janabi, M.; Lesman-Segev, O. H.; Miller, Z. A.; Perry, D. C.; O’Neil, J. P.; Pham, J.; Rojas, J. C.; Rosen, H. J.; Seeley, W. W.; Tsai, R. M.; Miller, B. L.; Jagust, W. J.; Rabinovici, G. D. Prospective Longitudinal Atrophy in Alzheimer’s Disease Correlates with the Intensity and Topography of Baseline Tau-PET. Sci. Transl. Med. 2020, 12 (524), eaau5732. 10.1126/scitranslmed.aau5732.

(17) Therriault, J.; Benedet, A. L.; Pascoal, T. A.; Mathotaarachchi, S.; Chamoun, M.; Savard, M.; Thomas, E.; Kang, M. S.; Lussier, F.; Tissot, C.; Parsons, M.; Qureshi, M. N. I.; Vitali, P.; Massarweh, G.; Soucy, J.-P.; Rej, S.; Saha-Chaudhuri, P.; Gauthier, S.; Rosa-Neto, P. Association of Apolipoprotein E Ε4 With Medial Temporal Tau Independent of Amyloid-β. JAMA Neurol 2020, 77 (4), 470. 10.1001/jamaneurol.2019.4421.

(18) Steward, A.; Biel, D.; Dewenter, A.; Roemer, S.; Wagner, F.; Dehsarvi, A.; Rathore, S.; Otero Svaldi, D.; Higgins, I.; Brendel, M.; Dichgans, M.; Shcherbinin, S.; Ewers, M.; Franzmeier, N. ApoE4 and Connectivity-Mediated Spreading of Tau Pathology at Lower Amyloid Levels. JAMA Neurol 2023, 80 (12), 1295. 10.1001/jamaneurol.2023.4038.

(19) Koutsodendris, N.; Blumenfeld, J.; Agrawal, A.; Traglia, M.; Grone, B.; Zilberter, M.; Yip, O.; Rao, A.; Nelson, M. R.; Hao, Y.; Thomas, R.; Yoon, S. Y.; Arriola, P.; Huang, Y. Neuronal APOE4 Removal Protects against Tau-Mediated Gliosis, Neurodegeneration and Myelin Deficits. Nat Aging 2023, 3 (3), 275–296. 10.1038/s43587-023-00368-3.

(20) Lennol, M. P.; Bordier, C.; Kamelher, L.; Ulrich, J. D.; Holtzman, D. M.; Gratuze, M. ApoE-Calypse Tau: ApoE–Tau Synergy in Alzheimer’s Disease. Journal of Experimental Medicine 2025, 222 (10), e20250965. 10.1084/jem.20250965.

(21) Nelson, M. R.; Liu, P.; Agrawal, A.; Yip, O.; Blumenfeld, J.; Traglia, M.; Kim, M. J.; Koutsodendris, N.; Rao, A.; Grone, B.; Hao, Y.; Yoon, S. Y.; Xu, Q.; De Leon, S.; Choenyi, T.; Thomas, R.; Lopera, F.; Quiroz, Y. T.; Arboleda-Velasquez, J. F.; Reiman, E. M.; Mahley, R. W.; Huang, Y. The APOE-R136S Mutation Protects against APOE4-Driven Tau Pathology, Neurodegeneration and Neuroinflammation. Nat Neurosci 2023, 26 (12), 2104–2121. 10.1038/s41593-023-01480-8.

(22) Shi, Y. The Role of Apolipoprotein E in Regulating Tau Pathogenesis and Neurodegeneration in a Tauopathy Mouse Model, Washington University in St. Louis, 2025. 10.7936/AV08-4126.

(23) Namba, Y.; Tomonaga, M.; Kawasaki, H.; Otomo, E.; Ikeda, K. Apolipoprotein E Immunoreactivity in Cerebral Amyloid Deposits and Neurofibrillary Tangles in Alzheimer’s Disease and Kuru Plaque Amyloid in Creutzfeldt-Jakob Disease. Brain Research 1991, 541 (1), 163–166. 10.1016/0006-8993(91)91092-F.

(24) Rohn, T. T.; Catlin, L. W.; Coonse, K. G.; Habig, J. W. Identification of an Amino -Terminal Fragment of Apolipoprotein E4 That Localizes to Neurofibrillary Tangles of the Alzheimer’s Disease Brain. Brain Research 2012, 1475, 106–115. 10.1016/j.brainres.2012.08.003.

(25) Agosta, F.; Vossel, K. A.; Miller, B. L.; Migliaccio, R.; Bonasera, S. J.; Filippi, M.; Boxer, A. L.; Karydas, A.; Possin, K. L.; Gorno-Tempini, M. L. Apolipoprotein E Ε4 Is Associated with Disease-Specific Effects on Brain Atrophy in Alzheimer’s Disease and Frontotemporal Dementia. Proc. Natl. Acad. Sci. U.S.A. 2009, 106 (6), 2018–2022. 10.1073/pnas.0812697106.

(26) Mishra, A.; Ferrari, R.; Heutink, P.; Hardy, J.; Pijnenburg, Y.; Posthuma, D. Gene-Based Association Studies Report Genetic Links for Clinical Subtypes of Frontotemporal Dementia. Brain 2017, 140 (5), 1437–1446. 10.1093/brain/awx066.

(27) Engelborghs, S.; Dermaut, B.; Marien, P.; Symons, A.; Vloeberghs, E.; Maertens, K.; Somers, N.; Goeman, J.; Rademakers, R.; Vandenbroeck, M. Dose Dependent Effect of APOE Ɛ4 on Behavioral Symptoms in Frontal Lobe Dementia. Neurobiology of Aging 2006, 27 (2), 285–292. 10.1016/j.neurobiolaging.2005.02.005.

(28) Koriath, C.; Lashley, T.; Taylor, W.; Druyeh, R.; Dimitriadis, A.; Denning, N.; Williams, J.; Warren, J. D.; Fox, N. C.; Schott, J. M.; Rowe, J. B.; Collinge, J.; Rohrer, J. D.; Mead, S. ApoE4 Lowers Age at Onset in Patients with Frontotemporal Dementia and Tauopathy Independent of Amyloid-β Copathology. Alz & Dem Diag Ass & Dis Mo 2019, 11 (1), 277–280. 10.1016/j.dadm.2019.01.010.

(29) Boyko, S.; Surewicz, W. K. Tau Liquid–Liquid Phase Separation in Neurodegenerative Diseases. Trends in Cell Biology 2022, 32 (7), 611–623. 10.1016/j.tcb.2022.01.011.

(30) Wegmann, S.; Eftekharzadeh, B.; Tepper, K.; Zoltowska, K. M.; Bennett, R. E.; Dujardin, S.; Laskowski, P. R.; MacKenzie, D.; Kamath, T.; Commins, C.; Vanderburg, C.; Roe, A. D.; Fan, Z.; Molliex, A. M.; Hernandez-Vega, A.; Muller, D.; Hyman, A. A.; Mandelkow, E.; Taylor, J. P.; Hyman, B. T. Tau Protein Liquid–Liquid Phase Separation Can Initiate Tau Aggregation. The EMBO Journal 2018, 37 (7), e98049. 10.15252/embj.201798049.

(31) Wen, J.; Hong, L.; Krainer, G.; Yao, Q.-Q.; Knowles, T. P. J.; Wu, S.; Perrett, S. Conformational Expansion of Tau in Condensates Promotes Irreversible Aggregation. J. Am. Chem. Soc. 2021, 143 (33), 13056–13064. 10.1021/jacs.1c03078.

(32) Zhang, X.; Vigers, M.; McCarty, J.; Rauch, J. N.; Fredrickson, G. H.; Wilson, M. Z.; Shea, J.-E.; Han, S.; Kosik, K. S. The Proline-Rich Domain Promotes Tau Liquid–Liquid Phase Separation in Cells. Journal of Cell Biology 2020, 219 (11), e202006054. 10.1083/jcb.202006054.

(33) Soeda, Y.; Yoshimura, H.; Bannai, H.; Koike, R.; Shiiba, I.; Takashima, A. Intracellular Tau Fragment Droplets Serve as Seeds for Tau Fibrils. Structure 2024, 32 (10), 1793–1807.e6. 10.1016/j.str.2024.06.018.

(34) Lin, Y.; Fichou, Y.; Longhini, A. P.; Llanes, L. C.; Yin, P.; Bazan, G. C.; Kosik, K. S.; Han, S. Liquid -Liquid Phase Separation of Tau Driven by Hydrophobic Interaction Facilitates Fibrillization of Tau. Journal of Molecular Biology 2021, 433 (2), 166731. 10.1016/j.jmb.2020.166731.

(35) Zhang, X.; Lin, Y.; Eschmann, N. A.; Zhou, H.; Rauch, J. N.; Hernandez, I.; Guzman, E.; Kosik, K. S.; Han, S. RNA Stores Tau Reversibly in Complex Coacervates. PLoS Biol 2017, 15 (7), e2002183. 10.1371/journal.pbio.2002183.

(36) Hernández-Vega, A.; Braun, M.; Scharrel, L.; Jahnel, M.; Wegmann, S.; Hyman, B. T.; Alberti, S.; Diez, S.; Hyman, A. A. Local Nucleation of Microtubule Bundles through Tubulin Concentration into a Condensed Tau Phase. Cell Reports 2017, 20 (10), 2304–2312. 10.1016/j.celrep.2017.08.042.

(37) Longfield, S. F.; Mollazade, M.; Wallis, T. P.; Gormal, R. S.; Joensuu, M.; Wark, J. R.; van Waardenberg, A. J.; Small, C.; Graham, M. E.; Meunier, F. A.; Martínez-Mármol, R. Tau Forms Synaptic Nano-Biomolecular Condensates Controlling the Dynamic Clustering of Recycling Synaptic Vesicles. Nat Commun 2023, 14 (1), 7277. 10.1038/s41467-023-43130-4.

(38) Boyko, S.; Surewicz, K.; Surewicz, W. K. Regulatory Mechanisms of Tau Protein Fibrillation under the Conditions of Liquid–Liquid Phase Separation. Proc. Natl. Acad. Sci. U.S.A. 2020, 117 (50), 31882–31890. 10.1073/pnas.2012460117.

(39) Kanaan, N. M.; Hamel, C.; Grabinski, T.; Combs, B. Liquid-Liquid Phase Separation Induces Pathogenic Tau Conformations in Vitro. Nat Commun 2020, 11 (1), 2809. 10.1038/s41467-020-16580-3.

(40) Michaels, T. C. T.; Qian, D.; Šarić, A.; Vendruscolo, M.; Linse, S.; Knowles, T. P. J. Amyloid Formation as a Protein Phase Transition. Nat Rev Phys 2023, 5 (7), 379–397. 10.1038/s42254-023-00598-9.

(41) Lin, Y.; Fichou, Y.; Longhini, A. P.; Llanes, L. C.; Yin, P.; Bazan, G. C.; Kosik, K. S.; Han, S. Liquid -Liquid Phase Separation of Tau Driven by Hydrophobic Interaction Facilitates Fibrillization of Tau. Journal of Molecular Biology 2021, 433 (2), 166731. 10.1016/j.jmb.2020.166731.

(42) Sahu, M. S. R.; Wei, J.; Weitz, D. A.; Knowles, T. P. J.; Garai, K. Liquid-Solid Coexistence at Single Fibril Resolution during Tau Condensate Ageing. Biophysics February 8, 2025. 10.1101/2025.02.06.636870.

(43) Wang, Y.; Mandelkow, E. Tau in Physiology and Pathology. Nat Rev Neurosci 2016, 17 (1), 22–35. 10.1038/nrn.2015.1.

(44) Kang, M.; Day, C. A.; Kenworthy, A. K.; DiBenedetto, E. Simplified Equation to Extract Diffusion Coefficients from Confocal FRAP Data. Traffic 2012, 13 (12), 1589–1600. 10.1111/tra.12008.

(45) Linsenmeier, M.; Faltova, L.; Morelli, C.; Capasso Palmiero, U.; Seiffert, C.; Küffner, A. M.; Pinotsi, D.; Zhou, J.; Mezzenga, R.; Arosio, P. The Interface of Condensates of the hnRNPA1 Low-Complexity Domain Promotes Formation of Amyloid Fibrils. Nat. Chem. 2023, 15 (10), 1340–1349. 10.1038/s41557-023-01289-9.

(46) Choi, C.-H.; Lee, D. S. W.; Sanders, D. W.; Brangwynne, C. P. Condensate Interfaces Can Accelerate Protein Aggregation. Biophysical Journal 2024, 123 (11), 1404–1413. 10.1016/j.bpj.2023.10.009.

(47) Dasadhikari, S.; Ghosh, S.; Pal, S.; Knowles, T. P. J.; Garai, K. A Single Fibril Study Reveals That ApoE Inhibits the Elongation of Aβ42 Fibrils in an Isoform-Dependent Manner. Commun Chem 2025, 8 (1), 133. 10.1038/s42004-025-01524-z.

(48) Hauser, P. S.; Narayanaswami, V.; Ryan, R. O. Apolipoprotein E: From Lipid Transport to Neurobiology. Progress in Lipid Research 2011, 50 (1), 62–74. 10.1016/j.plipres.2010.09.001.

(49) Frieden, C.; Wang, H.; Ho, C. M. W. A Mechanism for Lipid Binding to apoE and the Role of Intrinsically Disordered Regions Coupled to Domain–Domain Interactions. Proc. Natl. Acad. Sci. U.S.A. 2017, 114 (24), 6292–6297. 10.1073/pnas.1705080114.

(50) Garai, K.; Baban, B.; Frieden, C. Dissociation of Apolipoprotein E Oligomers to Monomer Is Required for High-Affinity Binding to Phospholipid Vesicles. Biochemistry 2011, 50 (13), 2550–2558. 10.1021/bi1020106.

(51) Lindner, K.; Beckenbauer, K.; Van Ek, L. C.; Titeca, K.; De Leeuw, S. M.; Awwad, K.; Hanke, F.; Korepanova, A. V.; Rybin, V.; Van Der Kam, E. L.; Mohler, E. G.; Tackenberg, C.; Lakics, V.; Gavin, A.-C. Isoform- and Cell-State-Specific Lipidation of ApoE in Astrocytes. Cell Reports 2022, 38 (9), 110435. 10.1016/j.celrep.2022.110435.

(52) Heinsinger, N. M.; Gachechiladze, M. A.; Rebeck, G. W. Apolipoprotein E Genotype Affects Size of ApoE Complexes in Cerebrospinal Fluid. J Neuropathol Exp Neurol 2016, 75 (10), 918–924. 10.1093/jnen/nlw067.

(53) Morikawa, M.; Fryer, J. D.; Sullivan, P. M.; Christopher, E. A.; Wahrle, S. E.; DeMattos, R. B.; O’Dell, M. A.; Fagan, A. M.; Lashuel, H. A.; Walz, T.; Asai, K.; Holtzman, D. M. Production and Characterization of Astrocyte-Derived Human Apolipoprotein E Isoforms from Immortalized Astrocytes and Their Interactions with Amyloid-β. Neurobiology of Disease 2005, 19 (1–2), 66–76. 10.1016/j.nbd.2004.11.005.

(54) Dasadhikari, S.; Ghosh, S.; Pal, S.; Knowles, T. P. J.; Garai, K. A Single Fibril Study Reveals That ApoE Inhibits the Elongation of Aβ42 Fibrils in an Isoform-Dependent Manner. Commun Chem 2025, 8 (1), 133. 10.1038/s42004-025-01524-z.

(55) Nguyen, D.; Dhanasekaran, P.; Nickel, M.; Nakatani, R.; Saito, H.; Phillips, M. C.; Lund -Katz, S. Molecular Basis for the Differences in Lipid and Lipoprotein Binding Properties of Human Apolipoproteins E3 and E4. Biochemistry 2010, 49 (51), 10881–10889. 10.1021/bi1017655.

(56) Cruchaga, C.; Kauwe, J. S. K.; Harari, O.; Jin, S. C.; Cai, Y.; Karch, C. M.; Benitez, B. A.; Jeng, A. T.; Skorupa, T.; Carrell, D.; Bertelsen, S.; Bailey, M.; McKean, D.; Shulman, J. M.; De Jager, P. L.; Chibnik, L.; Bennett, D. A.; Arnold, S. E.; Harold, D.; Sims, R.; Gerrish, A.; Williams, J.; Van Deerlin, V. M.; Lee, V. M.-Y.; Shaw, L. M.; Trojanowski, J. Q.; Haines, J. L.; Mayeux, R.; Pericak-Vance, M. A.; Farrer, L. A.; Schellenberg, G. D.; Peskind, E. R.; Galasko, D.; Fagan, A. M.; Holtzman, D. M.; Morris, J. C.; Goate, A. M. GWAS of Cerebrospinal Fluid Tau Levels Identifies Risk Variants for Alzheimer’s Disease. Neuron 2013, 78 (2), 256–268. 10.1016/j.neuron.2013.02.026.

(57) Alzheimer’s Disease Neuroimaging Initiative (ADNI); The Alzheimer Disease Genetic Consortium (ADGC); Deming, Y.; Li, Z.; Kapoor, M.; Harari, O.; Del-Aguila, J. L.; Black, K.; Carrell, D.; Cai, Y.; Fernandez, M. V.; Budde, J.; Ma, S.; Saef, B.; Howells, B.; Huang, K.; Bertelsen, S.; Fagan, A. M.; Holtzman, D. M.; Morris, J. C.; Kim, S.; Saykin, A. J.; De Jager, P. L.; Albert, M.; Moghekar, A.; O’Brien, R.; Riemenschneider, M.; Petersen, R. C.; Blennow, K.; Zetterberg, H.; Minthon, L.; Van Deerlin, V. M.; Lee, V. M.-Y.; Shaw, L. M.; Trojanowski, J. Q.; Schellenberg, G.; Haines, J. L.; Mayeux, R.; Pericak-Vance, M. A.; Farrer, L. A.; Peskind, E. R.; Li, G.; Di Narzo, A. F.; Kauwe, J. S. K.; Goate, A. M.; Cruchaga, C. Genome-Wide Association Study Identifies Four Novel Loci Associated with Alzheimer’s Endophenotypes and Disease Modifiers. Acta Neuropathol 2017, 133 (5), 839–856. 10.1007/s00401-017-1685-y.

(58) Strittmatter, W. J.; Saunders, A. M.; Goedert, M.; Weisgraber, K. H.; Dong, L. M.; Jakes, R.; Huang, D. Y.; Pericak-Vance, M.; Schmechel, D.; Roses, A. D. Isoform-Specific Interactions of Apolipoprotein E with Microtubule-Associated Protein Tau: Implications for Alzheimer Disease. Proc. Natl. Acad. Sci. U.S.A. 1994, 91 (23), 11183–11186. 10.1073/pnas.91.23.11183.

(59) Fleming, L. M.; Weisgraber, K. H.; Strittmatter, W. J.; Troncoso, J. C.; Johnson, G. V. W. Differential Binding of Apolipoprotein E Isoforms to Tau and Other Cytoskeletal Proteins. Experimental Neurology 1996, 138 (2), 252–260. 10.1006/exnr.1996.0064.

(60) Brecht, W. J.; Harris, F. M.; Chang, S.; Tesseur, I.; Yu, G.-Q.; Xu, Q.; Dee Fish, J.; Wyss-Coray, T.; Buttini, M.; Mucke, L.; Mahley, R. W.; Huang, Y. Neuron-Specific Apolipoprotein E4 Proteolysis Is Associated with Increased Tau Phosphorylation in Brains of Transgenic Mice. J. Neurosci. 2004, 24 (10), 2527–2534. 10.1523/JNEUROSCI.4315-03.2004.

(61) Harris, F. M.; Brecht, W. J.; Xu, Q.; Mahley, R. W.; Huang, Y. Increased Tau Phosphorylation in Apolipoprotein E4 Transgenic Mice Is Associated with Activation of Extracellular Signal -Regulated Kinase. Journal of Biological Chemistry 2004, 279 (43), 44795–44801. 10.1074/jbc.M408127200.

(62) Wang, C.; Xiong, M.; Gratuze, M.; Bao, X.; Shi, Y.; Andhey, P. S.; Manis, M.; Schroeder, C.; Yin, Z.; Madore, C.; Butovsky, O.; Artyomov, M.; Ulrich, J. D.; Holtzman, D. M. Selective Removal of Astrocytic APOE4 Strongly Protects against Tau-Mediated Neurodegeneration and Decreases Synaptic Phagocytosis by Microglia. Neuron 2021, 109 (10), 1657–1674.e7. 10.1016/j.neuron.2021.03.024.

(63) Litvinchuk, A.; Huynh, T. V.; Shi, Y.; Jackson, R. J.; Finn, M. B.; Manis, M.; Francis, C. M.; Tran, A. C.; Sullivan, P. M.; Ulrich, J. D.; Hyman, B. T.; Cole, T.; Holtzman, D. M. Apolipoprotein E4 Reduction with Antisense Oligonucleotides Decreases Neurodegeneration in a Tauopathy Model. Annals of Neurology 2021, 89 (5), 952–966. 10.1002/ana.26043.

(64) Yamada, K.; Cirrito, J. R.; Stewart, F. R.; Jiang, H.; Finn, M. B.; Holmes, B. B.; Binder, L. I.; Mandelkow, E.-M.; Diamond, M. I.; Lee, V. M.-Y.; Holtzman, D. M. *In Vivo* Microdialysis Reveals Age-Dependent Decrease of Brain Interstitial Fluid Tau Levels in P301S Human Tau Transgenic Mice. J. Neurosci. 2011, 31 (37), 13110–13117. 10.1523/JNEUROSCI.2569-11.2011.

(65) Clavaguera, F.; Hench, J.; Goedert, M.; Tolnay, M. Invited Review: Prion-like Transmission and Spreading of Tau Pathology. Neuropathology Appl Neurobio 2015, 41 (1), 47–58. 10.1111/nan.12197.

(66) Montal, V.; Diez, I.; Kim, C.-M.; Orwig, W.; Bueichekú, E.; Gutiérrez-Zúñiga, R.; Bejanin, A.; Pegueroles, J.; Dols-Icardo, O.; Vannini, P.; El-Fakhri, G.; Johnson, K. A.; Sperling, R. A.; Fortea, J.; Sepulcre, J. Network Tau Spreading Is Vulnerable to the Expression Gradients of *APOE* and Glutamatergic-Related Genes. Sci. Transl. Med. 2022, 14 (655), eabn7273. 10.1126/scitranslmed.abn7273.

(67) Chen, G.; Wang, M.; Zhang, Z.; Hong, D. K.; Ahn, E. H.; Liu, X.; Kang, S. S.; Ye, K. ApoE3 R136S Binds to Tau and Blocks Its Propagation, Suppressing Neurodegeneration in Mice with Alzheimer’s Disease. Neuron 2025, 113 (5), 719–736.e5. 10.1016/j.neuron.2024.12.015.

(68) Shi, Y.; Andhey, P. S.; Ising, C.; Wang, K.; Snipes, L. L.; Boyer, K.; Lawson, S.; Yamada, K.; Qin, W.; Manis, M.; Serrano, J. R.; Benitez, B. A.; Schmidt, R. E.; Artyomov, M.; Ulrich, J. D.; Holtzman, D. M. Overexpressing Low-Density Lipoprotein Receptor Reduces Tau-Associated Neurodegeneration in Relation to apoE-Linked Mechanisms. Neuron 2021, 109 (15), 2413–2426.e7. 10.1016/j.neuron.2021.05.034.

(69) Litvinchuk, A.; Suh, J. H.; Guo, J. L.; Lin, K.; Davis, S. S.; Bien-Ly, N.; Tycksen, E.; Tabor, G. T.; Remolina Serrano, J.; Manis, M.; Bao, X.; Lee, C.; Bosch, M.; Perez, E. J.; Yuede, C. M.; Cashikar, A. G.; Ulrich, J. D.; Di Paolo, G.; Holtzman, D. M. Amelioration of Tau and ApoE4-Linked Glial Lipid Accumulation and Neurodegeneration with an LXR Agonist. Neuron 2024, 112 (3), 384–403.e8. 10.1016/j.neuron.2023.10.023.

(70) Banani, S. F.; Lee, H. O.; Hyman, A. A.; Rosen, M. K. Biomolecular Condensates: Organizers of Cellular Biochemistry. Nat Rev Mol Cell Biol 2017, 18 (5), 285–298. 10.1038/nrm.2017.7.

(71) Rai, S. K.; Savastano, A.; Singh, P.; Mukhopadhyay, S.; Zweckstetter, M. Liquid–Liquid Phase Separation of Tau: From Molecular Biophysics to Physiology and Disease. Protein Science 2021, 30 (7), 1294–1314. 10.1002/pro.4093.

(72) Lin, Y.; Fichou, Y.; Longhini, A. P.; Llanes, L. C.; Yin, P.; Bazan, G. C.; Kosik, K. S.; Han, S. Liquid -Liquid Phase Separation of Tau Driven by Hydrophobic Interaction Facilitates Fibrillization of Tau. Journal of Molecular Biology 2021, 433 (2), 166731. 10.1016/j.jmb.2020.166731.

(73) Garai, K.; Mustafi, S. M.; Baban, B.; Frieden, C. Structural Differences between Apolipoprotein E3 and E4 as Measured by^19^ F NMR. Protein Science 2010, 19 (1), 66–74. 10.1002/pro.283.

(74) Garai, K.; Baban, B.; Frieden, C. Dissociation of Apolipoprotein E Oligomers to Monomer Is Required for High-Affinity Binding to Phospholipid Vesicles. Biochemistry 2011, 50 (13), 2550–2558. 10.1021/bi1020106.

(75) Ghosh, S.; Sil, T. B.; Dolai, S.; Garai, K. High-affinity Multivalent Interactions between Apolipoprotein E and the Oligomers of Amyloid-β. The FEBS Journal 2019, 286 (23), 4737–4753. 10.1111/febs.14988.

(76) Zimmermann, M. R.; Bera, S. C.; Meisl, G.; Dasadhikari, S.; Ghosh, S.; Linse, S.; Garai, K.; Knowles, T. P. J. Mechanism of Secondary Nucleation at the Single Fibril Level from Direct Observations of Aβ42 Aggregation. J. Am. Chem. Soc. 2021, 143 (40), 16621–16629. 10.1021/jacs.1c07228.

