## Supplementary Figures for "ApoE4 accelerates the condensate to amyloid transition of tau"

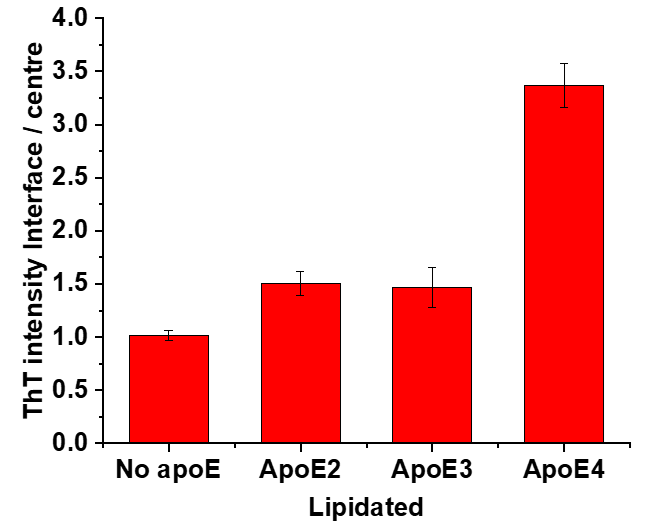

ThT F_ratio =_ F_interface/_ F_centre_

**Supplementary Fig 1:** Effect of lipidated apoE isoforms on the ratio of ThT fluorescence F_ratio_ F_interface_ / F_centre_ at the interface with respect to the centre of the condensates. The condensates were prepared using 20µM tau, 40µg/ml polyU, 5Um ThT and lipidated apoE isoforms. The samples were imaged using confocal microscopy. The representative images are shown in Fig 6D in the main manuscript.

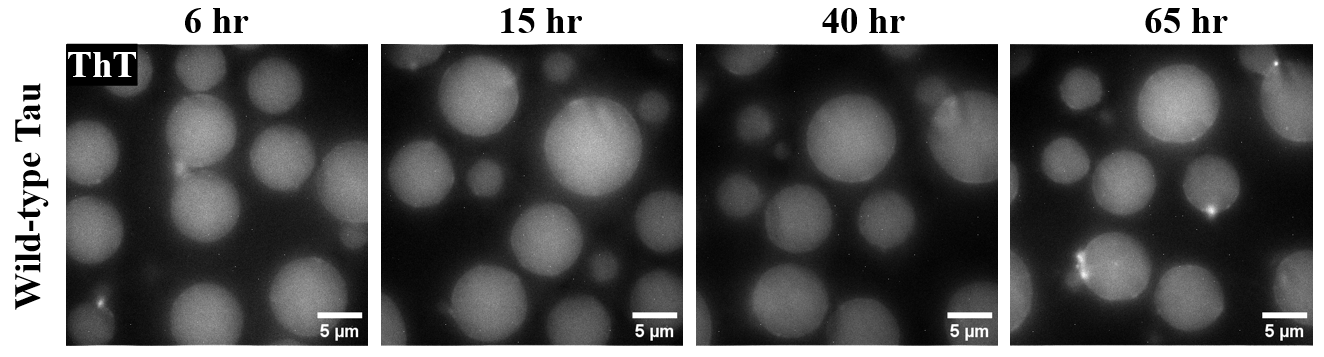

**Supplementary Fig 2:** Long term stability of the wild-type condensates. The condensates were imaged using ThT fluorescence till 65hr using fluorescence microscopy. Clearly, the condensates retain liquid nature till 65hr without transitioning to amyloid fibrils.
